# The CDK inhibitor SIAMESE targets both CDKA;1 and CDKB1 complexes to establish endoreplication in trichomes

**DOI:** 10.1101/2020.03.12.989079

**Authors:** Kai Wang, Ruth Ndathe, Narender Kumar, Elizabeth A. Zeringue, Naohiro Kato, John C. Larkin

## Abstract

Endoreplication, also known as endoreduplication, is a modified cell cycle in which DNA is replicated without subsequent cell division. Endoreplication plays important roles in both normal plant development and in stress responses. The *SIAMESE* (*SIM*) gene of Arabidopsis (*Arabidopsis thaliana*) encodes a cyclin-dependent kinase inhibitor that plays an central role in establishing endoreplication, and is the founding member of the *SIAMESE-RELATED* (*SMR*) family of plant-specific cyclin-dependent kinase inhibitors genes. However, there has been conflicting evidence regarding which specific cyclin/CDK complexes are inhibited by SIM in vivo. In this work, we use genetic evidence to show that SIM likely inhibits both CDKA;1- and CDKB1-containing CDK complexes in vivo to promote endoreplication in developing Arabidopsis trichomes. We also show that SIM interacts with CYCA2;3, a binding partner of CDKB1;1, via SIM Motif A, which we previously identified as a CDK-binding motif. In contrast, SIM Motif C, which has been indicated as a cyclin binding motif in other contexts, appears to be relatively unimportant for interaction between SIM and CYCA2;3. Together with earlier results, our work suggests that SIM and other SMRs likely have a multivalent interaction with CYC/CDK complexes.

**One sentence summary:** The cyclin-dependent kinase inhibitor SIAMESE (SIM) targets both CDKA;1 and CDKB1 complexes to establish endoreplication, and that SIM interacts with the cyclin CYCA2;3 via SIM Motif A.

## Introduction

Classically, the eukaryotic cell cycle is divided into four phases: G1, S, G2 and M, which are followed by cytokinesis. Cell cycle regulation depends in large part on specific cyclin and cyclin-dependent kinase (CDK) complexes that regulate both the G1→S and the G2→M transitions, which are the two major cell cycle checkpoints (Breuer, Ishida, & Sugimoto, 2010; Harashima, Dissmeyer, & Schnittger, 2013; Meijer & Murray, 2001; A. H. K. Roeder, 2012). In contrast to the mitotic cell cycle, cells can undergo endoreplication, an alternative cell cycle in which cells skip mitosis by inhibiting CDK/cyclin complex kinase activity, and continue to replicate their genomic DNA, resulting in increased DNA content in cells (De Veylder, Larkin, & Schnittger, 2011; Lee, Davidson, & Duronio, 2009).

Endoreplication, also known as endoreduplication, is common in higher plants, occurring during embryogenesis, tomato fruit development, and legume nitrogen-fixation root nodules, among other examples (Apri, Kromdijk, de Visser, de Gee, & Molenaar, 2014; Chevalier et al., 2014; De Veylder et al., 2011). Endoreplication is closely related to cell and organ growth, and is correlated with modification of cell walls (Bhosale, Maere, & De Veylder, 2019; Xu et al., 2016). Endoreplication can also occur in response to biotic or abiotic stress, such as response to light, temperature or drought stress, circadian clock disruption, pathogen defense, nematode feeding or DNA damage. (Adachi et al., 2011; De Veylder et al., 2011; Engler & Gheysen, 2013; Fung-Uceda et al., 2018; Hamdoun et al., 2016).

Arabidopsis (*Arabidopsis thaliana*) trichomes are a model for studying plant cell endoreplication. Wild type trichomes undergo endoreplication, reaching a DNA content of 16-32C (Walker, Oppenheimer, Concienne, & Larkin, 2000). A recessive mutant resulting from loss of *SIAMESE* (*SIM*) function was identified that disrupts endoreplication and allows mitosis to proceed, producing multicellular trichomes with a reduced DNA content per cell (Walker et al., 2000). In contrast, constitutively over-expressing *SIM* plants are small, with reduced leaves having enlarged epidermal cells that undergo increased endoreplication (Churchman et al., 2006). Thus, *SIM* negatively regulates mitosis and is required to initiate endoreplication to maintain *Arabidopsis* trichomes as single cells. In addition to their role in trichomes, *SIM* and its closest homolog *SIAMESE-RELATED1 (SMR1)* also play a role in initiating endoreplication in the root transition zone (Bhosale et al., 2018). The SIM protein can inhibit CDK activity in vitro, indicating that it likely functions as a CDK inhibitor in vivo (Kumar et al., 2015).

*SIM* was the first identified member of the plant-specific *SIAMESE-RELATED (SMR)* gene family, which is conserved in all land plant genomes. In *A*rabidopsis, the *SMRs* are represented by 17 genes. The biochemical function of *SMRs* appears to be largely equivalent, because several different *SMRs* can restore the unicellular trichome phenotype when expressed in the Arabidopsis *sim* mutant. Most significantly, an *SMR* from the bryophyte *Physcomitrella patens*, a distant relative of the angiosperms, can both suppress the *sim* multicellular trichome phenotype and inhibit CDK activity in vitro (Kumar et al., 2015). *SMRs* have only limited similarity to other types of CDK inhibitors (Churchman et al., 2006; Wang, Zhou, Bird, & Fowke, 2008). The SMR family is defined by three conserved protein motifs, termed motifs A, B and C (Kumar et al., 2015). Motif A has been implicated in interaction with CDKs (Kumar et al., 2018), and in a rice SMR, motif C is reported to be a cyclin-binding motif (Peres et al., 2007), although recent work has found that motif C of SIM is dispensable for suppression of mitosis in Arabidopsis trichomes (Kumar et al., 2018). Functions in plant growth and development have been identified for several of the *SMRs* including arrest of division and endoreplication in response to DNA damage (Yi et al., 2014), promotion of endoreplication in sepal giant cells (A. H. Roeder et al., 2010) and promotion of endoreplication in the root transition zone (Bhosale et al., 2018).

Several CDK/cyclin complexes have been implicated in promoting division and restricting endoreplication in *Arabidopsis*. Overexpression of *CYCD3;1* in trichomes causes mitosis instead of endoreplication, resulting in multicellular trichomes similar to those of *sim* mutants (Schnittger et al., 2002), while the triple mutant *cycd3;1 cycd3;2 cycd3;3* exhibits reduced division and increased endoreplication in leaves and petals (Dewitte et al., 2007). CYCD3;1 is known to activate CDKA;1 but not CDKB1;1 (Harashima & Schnittger, 2012; Nowack et al., 2012), suggesting that it is CYCD3;1/CDKA;1 complexes that are driving mitotic division at the expense of endoreplication in these instances. Another CYC/CDK complex implicated in promoting mitosis and suppressing endoreplication is the CYCA2;3/CDKB1;1 complex. Coexpression of *CDKB1;1* and *CYCA2;3* can suppress endoreplication in cotyledons (Boudolf et al., 2009). Also, overexpressing *CYCA2;3* including a mutated D-box that cannot mediate protein degradation further suppresses endoreplication in Arabidopsis (Imai et al., 2006).

Despite a great deal of work, it remains unclear which CYC/CDK complexes are the in vivo targets of inhibition by SIM to suppress mitosis and promote endoreplication. Initial studies implicated CYCD/CDKA;1 complexes as the primary interaction partners for SIM and other SMRs (Churchman et al., 2006; Peres et al., 2007). In contrast, an affinity-tagging proteomics study indicated that while most SMRs bound CYCD and CDKA;1, SIM and the closely related SMR1 protein bound to a CYCB and CDKB1;1, and not to CYCDs or CDKA;1 (Van Leene et al., 2010). More recently, interaction was detected between SIM and both CDKA;1 and CDKB1;1 in Arabidopsis protoplasts (Kumar et al., 2015), and another study found interaction of SMR1 with both CDKA;1 and CDKB1;1 in pulldown experiments from transgenic plant extracts (Dubois et al., 2018). Furthermore, genetic and biochemical studies show that both *CYCD3* function and *CDKB1* function are necessary for cell division in *sim* mutant trichomes, and that SIM can inhibit the kinase activity of both CDKA;1 and CDKB1;1 in vitro (Kumar et al., 2015).

The dichotomy between the naturally occurring endoreplication in wild-type trichomes and the contrasting mitotic division in *sim* mutant trichomes, combined with the other available mutants affecting CDKA;1 and CDKB1 complexes, provided us with a unique test system to test the effect of various cell cycle components on the balance between the endocycle and the mitotic cycle. In this study, we explored the roles of *CYCD3;1*, a presumed *CDKA;1* partner, and *CDKB1;1* in promoting cell division in Arabidopsis trichomes using a genetic approach, as well as exploring the protein interactions of SIM with known cyclin partners of CDKA;1 and CDKB1;1. Our results indicate that SIM likely inhibits both CDKA;1 and CDKB1 complexes in vivo. We also found that SIM interacts with CYCA2;3 primarily via Motif A. Conversely, SIM motif C is less important for interaction with CYCA2;3. Our results give new insights into the pathway by which SIM inhibits cyclin and CDK complexes to establish endoreplication in *Arabidopsis*.

## Results

### *CYCD3;1* overexpression can promote cell division in Arabidopsis trichomes in the absence of *CDKB1* function

Wild-type trichomes are unicellular (Fig 1A; Table 1), while the loss-of-function *sim* mutant has multicellular trichomes (Fig1B; Table 1). As previously reported by others (Schnittger et al., 2002), overexpression of *CYCD3;1* (*CYCD3;1^OE^*) in wild-type under control of the *GLABRA2* (*GL2*) trichome promoter resulted in cell division in trichomes (Fig1C; Table 1), and overexpression of *CYCD3;1* in *sim* mutants results in a greater degree of division (Fig1D; Table 1). To better understand the role of CYCD3;1 relative to CDKB1 in promoting cell division in trichomes, we took advantage of our earlier observation that *cdkb1;1 cdk1;2* double mutants (hereafter referred to as *cdkb1;1-2*) exclusively produce unicellular trichomes, and *sim cdkb1;1 cdkb1;2* triple mutants (hereafter referred to as *sim cdkb1;1-2*) exhibit only limited cell division in trichomes, mostly as rare clusters of two adjacent trichomes (Fig. 1E,F; Table 1; Kumar 2015). While these earlier observations show that the CDKB1s play a significant role in promoting cell division in *sim* mutant trichomes, it also afforded us with an opportunity in the present work to test whether *CYCD3;1^OE^* could promote cell division in the absence of CDKB1 function.

**Figure 1.**
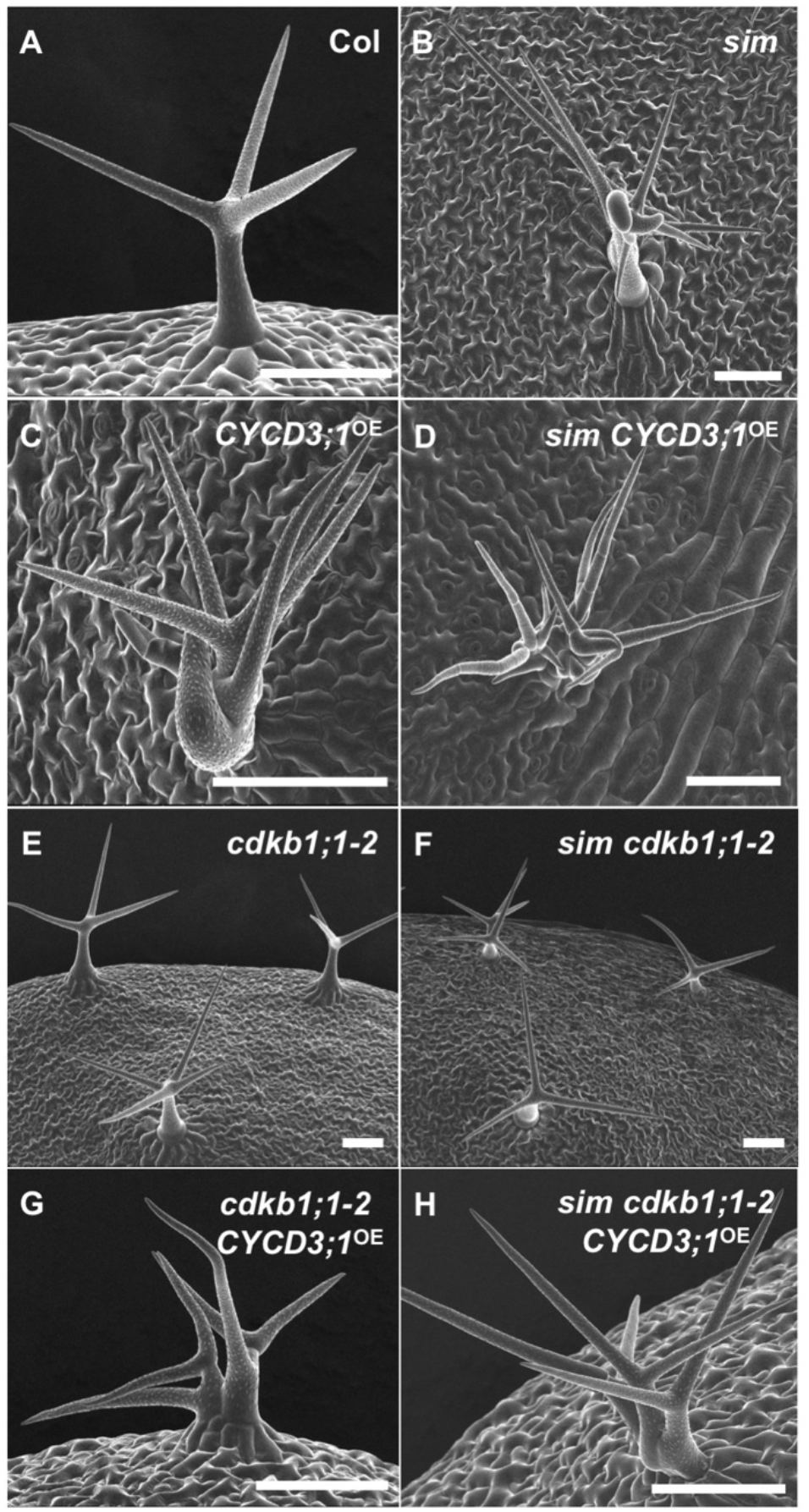
Cell division in Arabidopsis trichomes of various genotypes promoted by overexpression of *CYCD3;1* under control of the *GL2* promoter. Scanning electron micrographs of (A) Wild-type, (B) *sim*, (C) *CYCD3;1^OE^* in wild-type, (D) *sim CYCD3;1^OE^*, (E) *cdkb1;1-2*, (F) *sim cdkb1;1-2*, (G) *cdkb1;1-2 CYCD3;1^OE^*, (H) *sim cdkb1;1-2 CYCD3;1^OE^*. All scale bars 100μm.

**TABLE 1.** *CYCD3;1^oe^* can bypass the requirement for *CDKB1* to promote cell division in trichomes.

| Genotype | Homozygous line | Number of nuclei per TIS |
| --- | --- | --- |
| Col-0 | na | 1.02±0.25 |
| *sim* | na | 2.78±1.31 |
| *cdkb1;1-2* | na | 1.00±0.00 |
| *sim cdkb1;1-2* | na | 1.30±0.54 |
| Col-0 *CYCD3;1<sup>OE</sup>* | 2 | 1.40±0.76** |
|  | 4 | 4.06±2.87**** |
| *sim CYCD3;1<sup>OE</sup>* | 18 | 23.36±9.51**** |
|  | 19 | 11.48±6.10**** |
| *cdkb1;1-2 CYCD3;1<sup>OE</sup>* | 9 | 1.60±0.81**** |
|  | 10 | 1.22±0.42*** |
| *sim cdkb1;1-2 CYCD3;1<sup>OE</sup>* | 5 | 2.20±0.95**** |
|  | 10 | 2.14±0.76**** |

The phenotype of trichomes in control genotypes and homozygous single-insert *CYCD3;1<sup>OE</sup>* transgenic lines was assessed by counting the number of DAPI-stained nuclei at each trichome initiation site (TIS) for 50 TIS per genotype. Each transgenic line was compared to the corresponding non-transgenic genotype (Col-0 *CYCD3;1<sup>OE</sup>* lines vs. Col-0 etc.) in a two-tailed t-test. Bonferroni-corrected p values are as follows:

Of twenty-one independent T2 lines of *cdkb1;1-2* transformed with the *CYCD3;1^OE^* construct, six lines showed increased trichome cell division relative to the original *cdkb1;1-2* parent line, which shows no cell division in trichomes (Table S1). From these lines, we derived two independent *CYCD3;1^OE^ cdkb1;1-2* homozygous single insert lines that showed significantly increased cell division relative to *cdkb1;1-2* (Fig. 1G, Table 1). Similarly, seven out of twenty-two independent T2 lines obtained from transformation of *sim cdkb1;1-2* with *CYCD3;1^OE^* showed an increase in cell division above that of the *sim cdkb1;1-2* parent line (Table S2), and two independent homozygous *CYCD3;1^OE^ sim cdkb1;1-2* single insert lines were derived that exhibit increased division in trichomes (Fig. 1H, Table 1). These results demonstrate that *CYCD3;1^OE^* can drive cell division in trichomes even in the absence of CDKB1 function.

### Other D-cyclins do not promote cell division in trichomes when expressed from the *GL2* promoter

Several other D-type cyclins have been implicated in promoting mitosis under certain circumstances (Kono et al., 2007; Qi & John, 2007; Sozzani et al., 2010). The ability of several of these other D-type cyclins to promote division was assessed by examining the phenotype of the transgenic lines in which *CYCD2;1*, *CYCD4;1* or *CYCD6;1* coding regions were expressed from the *GL2* promoter in a wild-type background.

Qi and John (2007) have reported that expression of wild-type *CYCD2;1* cDNA from the 35S promoter resulted in a cryptic splicing event excising exons 2 and 3, resulting in an mRNA encoding a truncated protein. To prevent this cryptic splicing in our work, we introduced silent mutations at these splice junctions, as they have described (Qi & John, 2007), in a construct that we named *CYCD2;1NS^OE^* (NS for Non-Spliceable). The closely related *CYCD4;1* gene has similar sequences at the junctions flanking exons 2 and 3 that could result in a similar cryptic splice removing these exons, and we introduced similar mutations into the *CYCD4;1NS^OE^* construct to eliminate the chance of cryptic splicing of this transgene. When these constructs were expressed in plants, transformants expressing the wild-type *CYCD2;1^OE^* coding region produced transcripts missing exons 2 and 3, as expected, while the *CYCD2;1NS^OE^* construct expressed transcripts of the correct size (Fig. 2A) and sequence. Both the *CYCD4;1^OE^* and the *CYCD4;1NS^OE^* constructs, as well as the CYCD6;1 construct, produced transcripts of the expected size (Fig. 2A) and sequence for full-length transcripts. Thus the wild-type *CYCD4;1* transcript does not undergo the cryptic splicing seen in with the wild-type *CYCD2;1* coding region, in spite of high sequence similarity between the two genes. Examination of >50 primary transformants, at least 12 T2 transgenic families, and a minimum of three homozygous single-insert lines for each of the constructs (*CYCD2;1NS^OE^*, *CYCD4;1^OE^* and *CYCD6;1^OE^*) revealed wild-type trichomes with no evidence of cell division in any case (Fig. 2B-E).

**Figure 2.**
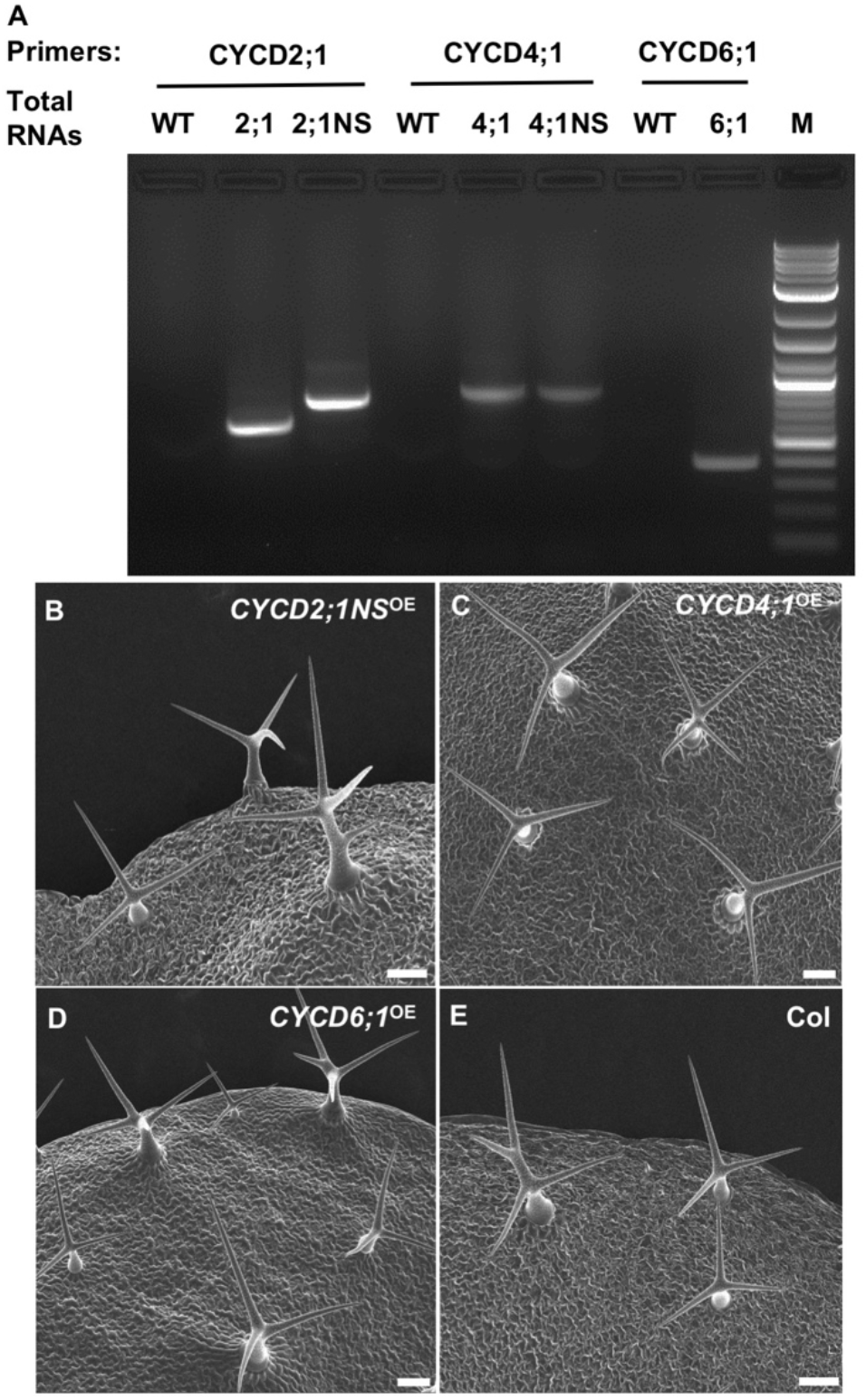
Expression of *CYCD2;1, CYCD4;1* or *CYCD6;1* from the *GL2* promoter does not cause cell division in Arabidopsis trichomes. (A) RT-PCR products from total RNA of wild-type (WT), a CYCD2;1^OE^ transformant, a CYCD2;1NS^OE^ transformant, a CYCD4;1^OE^ transformant, a CYCD4;1NS^OE^ transformant, and a CYCD6;1^OE^ transformant, amplified with the indicated primers. (B) CYCD2;1^OE^ trichomes expressing a coding region modified to prevent cryptic mis-splicing and predicted produce the correct protein product, (C) CYCD4;1^OE^ trichomes, (D) CYCD6;1^OE^ trichomes, (E) wild-type trichomes, Scale bars in (A), (B), (C) and (D) = 100μm.

### SIM interacts with CYCA2;3 but not CYCD3;1

CYCD3/CDKA;1 and CYCA2;3/CDKB1 complexes have both been implicated in promoting division and suppressing endoreplication (Boudolf et al., 2009; Schnittger et al., 2002). Previous work has shown that SIM can inhibit both CDKA;1 and CDKB1;1 complexes in vitro, and can bind to both types of CDK in Arabidopsis protoplasts. (Kumar et al., 2015). However, conflicting results have been reported in the literature regarding direct interactions of SIM with specific cyclins (Churchman et al., 2006; Van Leene et al., 2010). For this reason, we tested whether either CYCA2;3 or CYCD3;1 interacts with SIM using two different protein interaction assays.

The split luciferase complementation assay was adopted to test these interactions in Arabidopsis protoplasts (Fujikawa & Kato, 2007). For this assay, we tested the ability of SIM fused to the N-terminus of *Renilla reniformis* luciferase (Nluc:SIM) to interact with CYCA2;3 or CYCD3;1 fused to the C-terminal (Cluc) half of *Renilla reniformis* luciferase (CLuc:CYCA2;3 and CLuc:CYCD3;1, respectively). The interaction of histones H2A and H2B was used as a positive control. The interactions of SIM and the two cyclins with both the bZIP transcription factor PERIANTHIA (PAN) (Chuang, Running, Williams, & Meyerowitz, 1999) and with the histones were used as two independent negative controls. In this assay, SIM interacted with CYCA2;3 significantly more strongly than either protein interacted with the negative controls, while SIM showed no significant interaction with CYCD3;1 (Fig 3).

**Figure 3.**
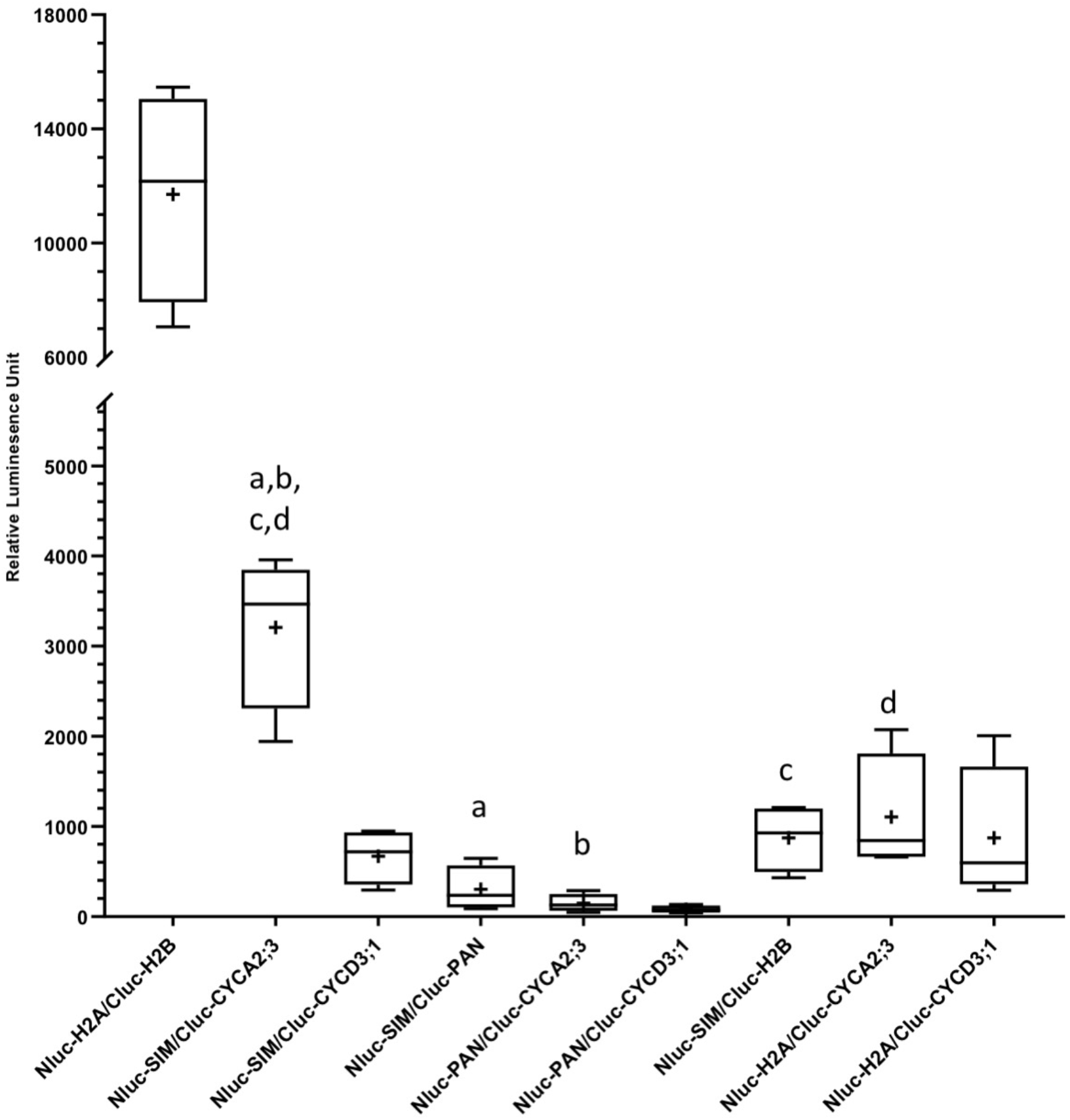
SIM interacts with CYCA2;3, but not with CYCD3;1, in a split-luciferase complementation assay. Nluc is the N-terminal portion of *Renilla reniformis* luciferase, and Cluc is the C-terminal portion of *Renilla reniformis* luciferase. Interaction of histone H2A with histone H2B was used as a positive control. Two different negative controls were used for SIM and each of the cyclins; first, interaction with the appropriate histone fusion, and second, interaction with fusions of the transcription factor PAN, NLuc:PAN and CLuc:PAN. The work presented is the result of four independent experimental trials, each of which included four technical replicates. In the box plots, central bar represents the median, box outline represents the first and third quartiles, whiskers extend to the maximum and minimum data point, and + represents the mean. Samples indicated with the same letter are significantly different (p<0.0001), based on a post hoc Tukey’s multiple comparison test, which was based on a two-way ANOVA on the two factors experimental trial and pairwise protein interactions (ANOVA summary statistics are given in Supplemental Table 5). Only relevant comparisons showing a significant difference are indicated. The comparisons of Nluc-SIM/Cluc-CYCD3;1 with Nluc-SIM/Cluc-PAN, Nluc-SIM/Cluc-H2B, Nluc-PAN/Cluc-CYCD3;1 and Nluc-H2A/Cluc-CYCD3;1 were not significant (p>0.70 in all four cases).

Interaction of SIM with CYCA2;3 was further tested with the yeast two-hybrid assay. CYCA2;3 and SIM were integrated into the vectors pASGW and pACTGW which contain the GAL4 DNA-binding domain and transcription activation domains, respectively. The resulting pASGW-CYCA2;3 and pACT-SIM constructs were introduced into yeast by cotransformation. SIM and CYCA2;3 showed clear interaction in a plate growth assay that is dependent on activation of the two selectable reporter genes, *HIS3* and *ADE2* (Fig. 4). The interaction of SIM and CYCA2;3 is also able to activate the *lacZ* reporter gene, further supporting this interaction (Supplemental Figure S1).

**Figure 4.**
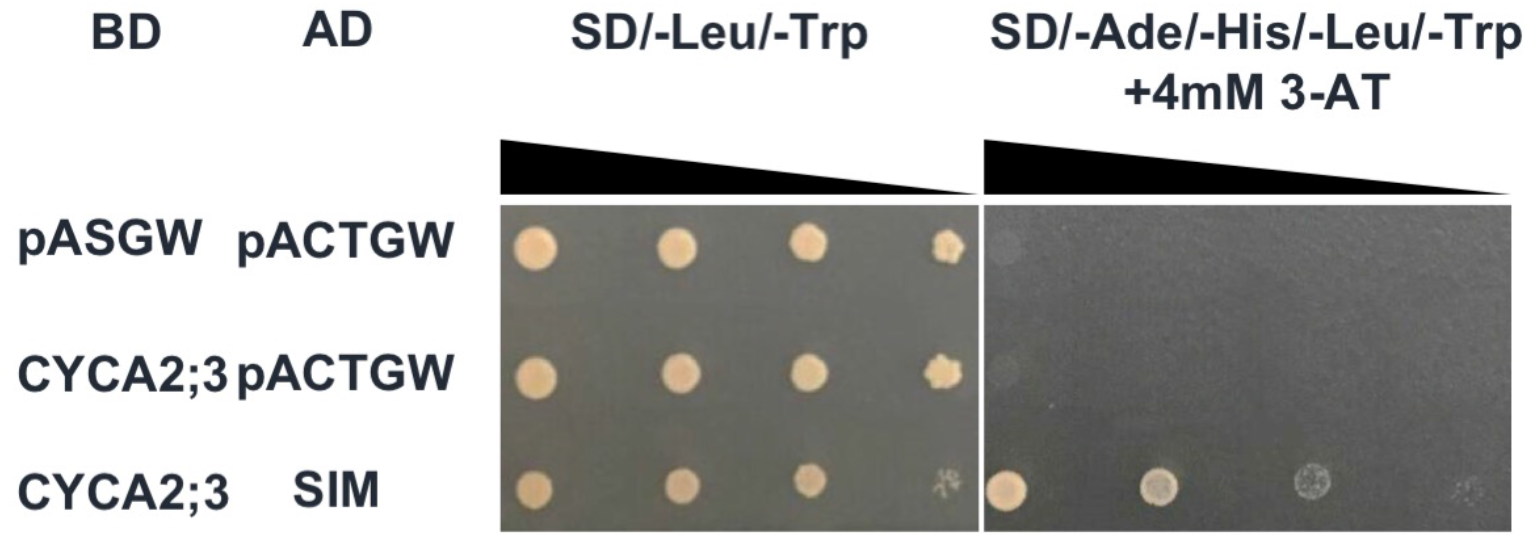
SIM interacts with CYCA2;3 in the yeast two-hybrid system. pASGW and pACTGW are empty vectors, including only binding domain or activation domain, respectively, and were used as negative controls. Yeast cultures of each genotype were diluted in the ratio of 1:10:100:1000 before spotting on plate made with the indicated drop-out media. 3-AT = 3-Amino-1,2,4-triazole.

### The conserved Thr-35 residue of SIM motif A is required for interaction between SIM and CYCA2;3

The predicted SIM protein sequence incudes three sequence motifs, motif A, motif B, and motif C (Fig. 5A), that define the SMR family, as well as two nuclear location signals (Kumar et al., 2018). To determine which of these motifs were essential for interaction with CYCA2;3, mutagenized versions of each motif (Fig. 5B) were tested for interaction with CYCA2;3 by yeast two-hybrid assay. Mutant constructs in which alanines replaced the three C-terminal residues of motif A (motA-3A), the four central residues of motif A (mot-4A) or the seven C-terminal residues of motif A (motA-7A) all showed interaction with CYCA2;3, while a mutant replacing all ten residues of motif A with alanines failed to show interaction (Fig. 5 B,C), suggesting that the N-terminal portion of this motif plays a role in the interaction between these two proteins. Mutation of motif B also eliminated the interaction (Fig. 5 B,C), possibly implicating this motif as well, though the same motif B mutant results in an unstable protein when expressed as a fluorescent protein fusion in Arabidopsis (Kumar et al., 2018). In contrast, a mutant in which the six key residues of motif C were replaced by alanines still interacted with CYCA2;3 (Fig. 5 B,C; Fig. S1).

**Figure 5.**
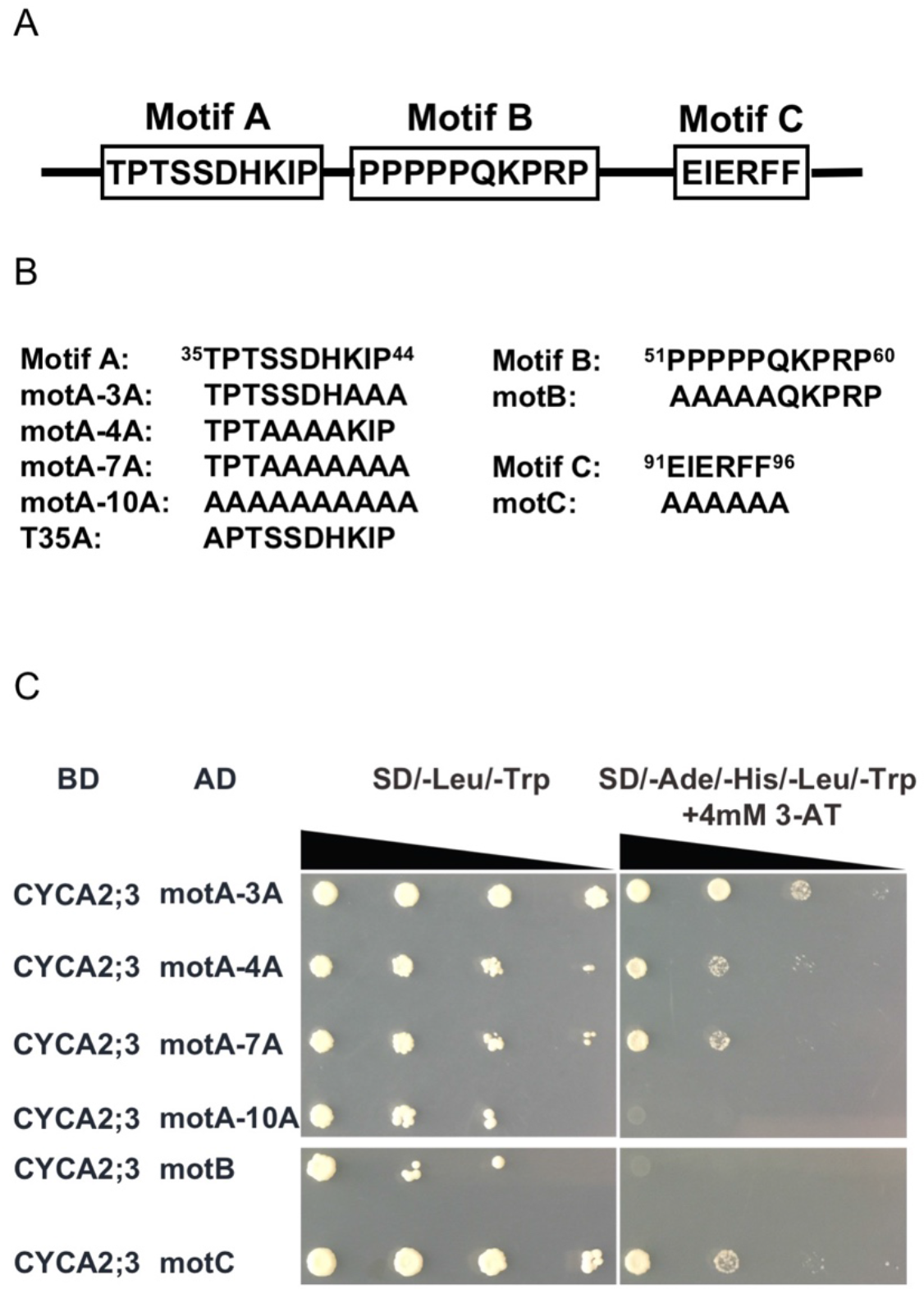
Identification of SIM protein sequence motifs responsible for interaction with CYCA2;3. (A) The sequence and arrangement of motifs A, B and C in the SIM protein. (B) Mutations of the SIM motifs that were tested for interaction. (C) Interaction of SIM motif mutants with CYCA2;3 in the yeast two-hybrid system. Yeast cultures of each genotype were diluted in the ratio of 1:10:100:1000 before spotting on plate made with the indicated drop-out media. 3-AT = 3-Amino-1,2,4-triazole.

Our recent work had identified residue T35, at the N-terminal end of motif A, as a key functional residue in SIM. When this residue is changed to alanine (T35A, Fig. 5B), the *SIM* gene cannot complement the *sim* mutant phenotype, while changing this residue to the phosphomimic amino acid aspartate (T35D) results in a functional gene that can fully complement *sim* (Kumar et al., 2018). The T35 residue is one of the three residues altered in the motA-10A mutant, but not in the other motif A mutants, and thus may play a significant role in the interaction with CYCA2;3. When tested, we found that both the T35A and T35D mutations eliminate the interaction (Fig. 6), indicating that T35 is a critical amino acid for the interaction between SIM and CYCA2;3.

**Figure 6.**
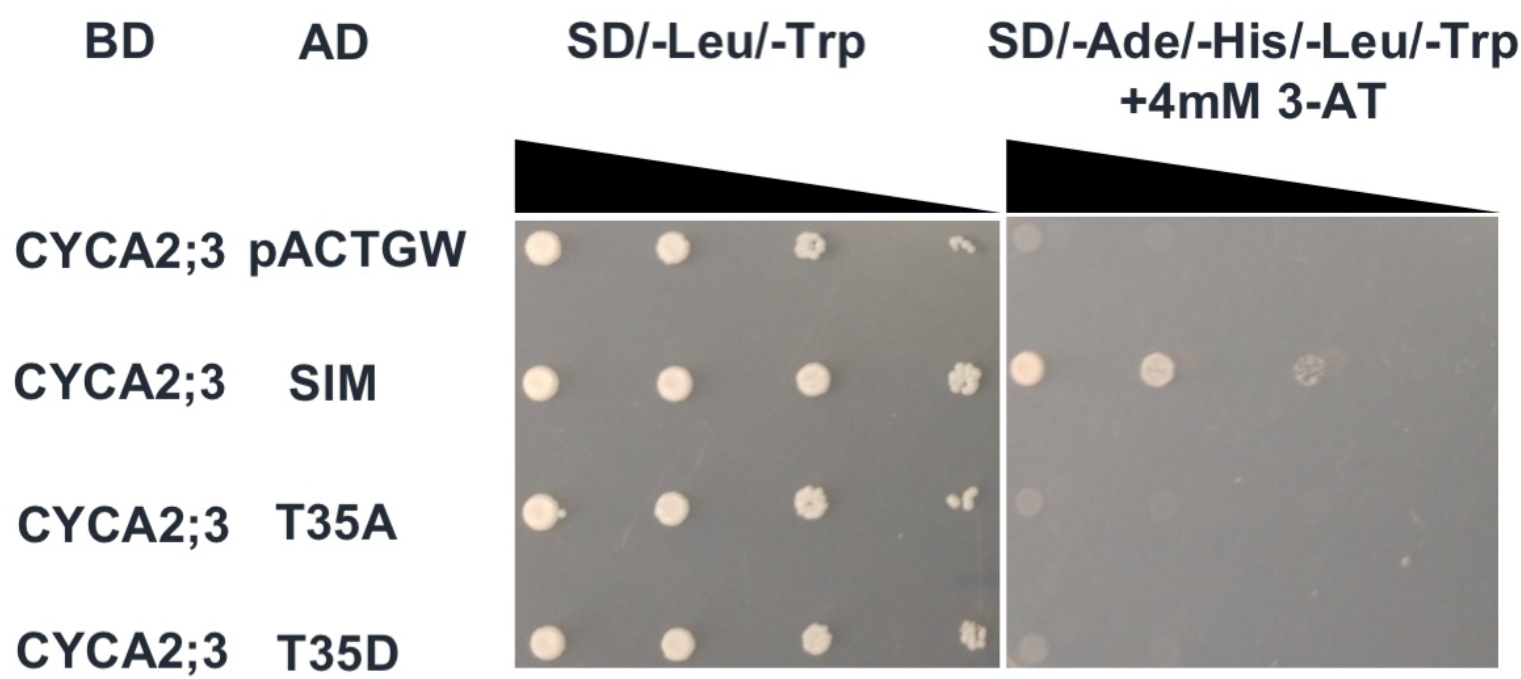
SIM residue T35 plays a crucial role in the interaction between SIM and CYCA2;3. Mutant *sim* constructs containing an alanine codon at position T35 (T35A) or an aspartate codon (T35D) were tested for interaction with CYCA2;3. Yeast cultures of each genotype were diluted in the ratio of 1:10:100:1000 before spotting on plate made with the indicated drop-out media. 3-AT = 3-Amino-1,2,4-triazole.

## Discussion

### Among D-cyclins, CYCD3;1 is uniquely able to promote cell division in developing trichomes

Although D-type cyclins are generally associated with promoting entry into S-phase, CYCD3;1 of Arabidopsis has in several instances been implicated in promoting mitosis and suppressing endoreplication, both from overexpression experiments and loss of function mutants (Dewitte et al., 2007; Schnittger et al., 2002). Consistent with a potential function in division, *CYCD3;1* is the only D-cyclin whose transcripts are expressed at their highest level in G2/M, rather than S-phase (Menges, de Jager, Gruissem, & Murray, 2005). Several other D-cyclins have also been associated with promoting cell proliferation. In transient expression experiments in *Nicotiana benthamiana*, co-expression of *CYCD4;1*, *CYCD4;2* or *CYCD5;1* with either *CDKA;1*or *CDKB1;1* induces ectopic cell divisions in tobacco epidermal cells (Boruc, Inze, & Russinova, 2010). Overexpression of correctly spliced *CYCD2;1* can promote cell division in both leaves and roots of Arabidopsis if expressed at a sufficient level (Qi & John, 2007). Loss-of-function *cycd4;1* or *cycd4;2* mutants have reduced proliferation in the stomatal lineage of the hypocotyl, while overexpression of either *CYCD4* paralog enhances cell division in this lineage (Kono et al., 2007). Similarly, *CYCD6;1* is specifically involved in the asymmetric division of the cortex/endodermal initial cells during root development (Sozzani et al., 2010). A naturally occurring *CYCD5;1* allele that exhibits increased gene expression relative to the standard Col-0 wild-type results in increased ploidy in leaves, and overexpression of *CYCD5;1* results in an increase in both cell proliferation and ploidy (Sterken et al., 2012).

Of the D-type cyclins that we tested, only *CYCD3;1* was found to be capable of promoting cell division in trichomes when expression of each cyclin was driven from the same *GL2* promoter, confirming the results of Schnittger et al. (2002) for CYCD4;1 and extending this to two other D-cyclins, *CYCD2;1* and *CYCD6;1* (Fig. 2). This result suggests that at least in trichomes, CYCD3;1 is relatively unique among D-type cyclins in its ability to promote cell division. It is now clear that individual cyclins can affect the target specificity of CYC/CDK complexes in plants (Harashima & Schnittger, 2012; Nowack et al., 2012). One possible explanation for our results is that CYCD3; 1-containing CDK complexes are more efficient than other CYCD/CDK complexes at phosphorylating specific target proteins, such as the MYB3R transcription factors, that are required for G2/M transcription and are activated by phosphorylation (Berckmans & De Veylder, 2009; Magyar, Bogre, & Ito, 2016). However, while we have confirmed expression of the correct transcripts at the mRNA level, we cannot rule out post-transcriptional effects on mRNA or protein stability, or post-translational protein modifications that may prevent CYCD2;1, CYCD4;1 or CYCD6;1 from functioning to promote mitosis in developing trichomes.

### CYCD3;1/CDKA;1 complexes and CDKB1 complexes act in parallel to promote cell division

Our results give insight into the order of function of CYC/CDK complexes in promoting mitosis, at least in the trichome system. Previous genetic evidence demonstrated that cell division of *sim* mutant trichomes is largely blocked in either the *sim cdkb1;1-2* triple mutant or the *sim cycd3;1 cycd3;2 cycd3;3* quadruple mutant (Kumar et al., 2015), indicating that both CDKB1 and CYCD3 are necessary for cell division in trichomes. CYCD3;1 is generally considered to form active CDK complexes only with CDKA;1, and not with CDKB1 (Berckmans & De Veylder, 2009; Harashima & Schnittger, 2012; Nowack et al., 2012; Van Leene et al., 2010), and the CDKB1 genes are transcribed in G2 and are thought to act exclusively in G2 and M (Boudolf et al., 2004; Menges et al., 2005). This might suggest a linear pathway in which CYCD3;1/CDKA acts first to activate CDKB1 function, and CDKB1 complexes are the kinase that is required for progression to mitosis.

Our demonstration that *CYCD3;1^oe^* can promote cell division in the *cdkb1;1-2* mutant rule out such a linear pathway, demonstrating that *CYCD3;1* can directly promote mitosis in the absence of CDKB1 function (Fig 1, G and H; Table 1, Table S1, Table S2). Additionally, it is noteworthy that in the homozygous T3 lines, as well as in the larger number of segregating T2 lines, the *CYCD3;1^OE^ sim cdkb1;1-2* lines exhibited more cell division per trichome initiation site than the *CYCD3;1^OE^ cdkb1;1-2* lines (Table S1, Table S2). While the phenotypic variability among individual transgenic lines indicates that quantitative comparisons between these two transgenic genotypes should be treated with caution, these observations provide evidence that SIM inhibits CYCD3;1-containing CDK complexes in vivo. Taken together, the results presented here, together with our previous results (Kumar et al., 2015), suggest that CYCD3/CDKA;1 and CYC/CDKB1 complexes act in parallel to promote cell proliferation, and that SIM can inhibit both types of CDK complex in its role promoting endoreplication.

### SIM can bind to CYCA2;3, a partner of CDKB1

It is by now clear that SIM can bind to and inhibit both CDKA;1 and CDKB;1 complexes, in vitro, and that binding to CDKs requires sequences in Motif A (Dubois et al., 2018; Kumar et al., 2018; Kumar et al., 2015; Van Leene et al., 2010). However, there is conflicting evidence about which cyclins SIM might bind to (Churchman et al., 2006; Van Leene et al., 2010), and no previous information on which sequences in SIM are involved in binding to cyclins. Our results indicate that SIM can bind to CYCA2;3 (Fig. 3 and 4), a previously verified partner of CDKB1;1 in Arabidopsis (Boudolf et al., 2009). *CYCA2;3* is expressed in trichomes, as well as in proliferating tissues. In trichomes, *CYCA2;3* is expressed after branching has been initiated, and acts to limit the degree of endoreplication (Imai et al., 2006). Thus, the interaction of SIM with CYCA2;3 that we have detected may be significant in wild-type trichomes, where SIM would be expected to counteract the endoreplication-inhibiting function of CYCA2;3/CDK complexes.

In contrast, we did not find evidence of direct binding between SIM and CYCD3;1 in our experiments, suggesting that SIM binds to CYCD3;1/CDKA;1 complexes primarily via interaction with CDKA;1. Previously, Churchman et al (2006) reported interaction of SIM with the closely related CYCD3;2 using acceptor-bleaching Förster resonance energy transfer (FRET). This method depends on extremely close proximity of the two fluorophores of less than 10nm (Xing, Wallmeroth, Berendzen, & Grefen, 2016), but does not require direct binding. It is possible that the reported FRET interaction signal with CYCD3;2 was due to interaction of both the cyclin and SIM fluorescent protein fusions with the ubiquitously expressed CDKA;1 that was present in the leaf cells, bringing the fluorophores close together.

### The conserved Thr-35 residue is important in SIM for interaction of SIM and CYCA2;3

Our results show that the binding of SIM and CYCA2;3 depends on the N-terminal end of MotifA and specifically on the conserved Thr-35 residue of motif A in SIM (Fig. 5C, Fig. 6). Motif A is also required for CDK binding of SIM. Thus Motif A may bind near the interface of the cyclin and the CDK. Our previous results showed that a mutation changing T35 to the non-phosphorylatable residue alanine (*T35A*) inactivates the biological function of *SIM*, while a mutant substituting the phosphomimic residue aspartate (*T35D*) was functional, suggesting that T35 is phosphorylated, and that this phosphorylation is required for function (Kumar et al., 2015). Interestingly, neither the T35A nor T35D mutant forms can bind to CYCA2;3 (Fig 6). Perhaps the phosphorylated form blocks cyclins from binding to their CDKs, thus preventing activation of the kinase.

Dubois et al (2018) have found that a threonine residue in a potential CDK phosphorylation site near the N-terminus of SMR1, the SMR most closely related to SIM, may play a role in ubiquitin-mediated proteolysis of SMR1. They have proposed that phosphorylation of this residue by CDKA;1 targets SMR1 for degradation. Based on this result, it has been suggested that CDKA;1 complexes inhibit function of both SMR1 and SIM, targeting them for degradation (Bhosale, Maere, & De Veylder, 2019). However, the potential CDK phosphorylation site in the *SMR1*-encoded polypeptide, T16, is not homologous to the T35 residue in Motif A of SIM, which in the *SMR1*-encoded protein is residue T43, and SIM does not have a threonine or serine at the position equivalent to T16 of SMR1. Our results here, combined with our previous results, clearly indicate that the T35 residue is required in a positive sense for SIM function, and that SIM is an inhibitor of CDKA;1 complexes both in vitro and in vivo.

### The role of Motif C

Our work here shows Motif C is not essential for interaction of SIM with CYCA2;3 (5C). Similarly, in previous work, we showed that Motif C is not necessary for in vivo function of *SIM* when the gene is overexpressed in developing trichomes (Kumar et al., 2018). Yet Motif C is conserved in the *SMR* family (Kumar et al., 2018), and has sequence similarity to a motif in the ICK/KRP family of cell cycle regulators that is required for ICK1/KRP1 interaction with CYCD3;1 (Churchman et al., 2006; De Veylder et al., 2002; Wang et al., 1998). Motif C is also required for interaction of the rice SMR *OsEL2* and with a rice D-cyclin (Peres et al., 2007). These results, together with the results we have presented here indicating the importance of Motif A in interaction with CYCA2;3, suggest that SMRs interact with cyclins via both Motif A and Motif C, and that the relative importance of these two motifs for cyclin binding likely differs among different cyclin/SMR pairs.

## Conclusion

We have shown that CYCD3;1, likely complexed with CDKA;1, and CDKB1 act in parallel to promote cell division in *sim* mutant trichomes. Our results also show that SIM interacts with CYCA2;3 via SIM Motif A, in contrast to earlier evidence implicating Motif C in cyclin-binding. These results also highlight the importance of the T35 residue in Motif A, which is the most conserved amino acid in the SMR family. Together with earlier results, our work suggests that SMRs likely have a multivalent interaction with CYC/CDK complexes that involves both Motifs A and C.

## Materials and Methods

### Plant growth and transgenic line generations

Plants were grown as previously described (Kumar et al., 2015). The *cdkb1;1 cdkb1;2* and *sim cdkb1;1cdkb1;2* homozygous mutant lines have described previously (Kumar et al., 2015). Primers used for reconfirming these genotypes are given in Supplemental Table S6. All transgenic lines including gene of interest in specific genetic background, like Col-0, *sim cdkb1;1 cdkb1;2* or *sim cdkb1;1 cdkb1;2* mutants were generated by homozygotes transformation by the floral dip method (Clough & Bent, 1998). Positive transformants were identified by Basta selection. T2 lines derived from different T1 transformants that showed segregation of 1:3 of Basta-sensitive to Basta-resistance were considered to be independent single-insertion transgenic lines.

### DNA Constructions

PCR primers used in all constructions are given in Supplemental Table 6. The plasmids pDONR221-CYCA2;3, ppHGGWA-His-GST-CYCD3;1 and pHMGWA His-MBP-CYCD6;1 were obtained from Dr. Hirofumi Harshima (U. of Strasbourg, France). The CYCA2;3, CYCD3;1 and CYCD6;1 coding sequence were PCR amplified from pDONR221-CYCA2;3, pHGGWA-His-GST-CYCD3;1 pHMGWA His-MBP-CYCD6;1, respectively by Phusion^®^ High-Fidelity PCR Kit (NEB) (Harashima & Schnittger, 2012)(Harashima,*et al*. 2012). CYCD2;1 and CYCD4;1 coding sequence were amplified from pDONR221 and pDEST14-CYCD4;1 templates. To introduce mutations into the potential cryptic splice sties in these two genes, an overlapping PCR reaction was conducted in which three separate products were amplified in the 1^st^ round of PCR from the appropriate coding region plasmid, then all products were combined, diluted with the ratio of 1 to 10, as the 2^nd^ round templates to produce the final CYCD2;1NS and CYCD4;1NS coding regions containing silent mutations in the potential cryptic splice junction sequences. All PCR products were purified with a QIAquick Gel Extraction Kit (Qiagen), and purified PCR fragments were inserted into the vector pENTR/D-TOPO using a pENTR Directional TOPO Cloning Kit (Life Technologies). The resulting entry clones were confirmed by sequencing. Error-free entry clones were integrated into the Gateway binary T-DNA destination vector pAMPAT-PROGL2 harboring the *GL2* promoter (Weinl et al., 2005) via LR Clonase reactions (Thermo Fisher). All mutant versions of *SIM* were described previously (Kumar et al., 2015). For yeast two-hybrid experiments, the wild-type and mutant *SIM* genes, CYCA2;3 and CYCD3;1-related entry clones were integrated via LR Clonase into either the pASGW-attR or the pACTGW-attR destination vectors (Nakayama, Kikuno, & Ohara, 2002)(Nakayama *et al*, 2002), which we refer to as pASGW or pACTGW in brief. All constructions were further confirmed by sequencing.

### RT-PCR

Total RNAs were extracted from two-week Arabidopsis seedlings following The RNeasy Plant Mini Kit protocol (QIAGEN), and 5μg total RNA was converted into cDNA with Oligo(dT)20 by SuperScript™ III First-Strand Synthesis System (Invitrogen). OneTaq^®^ DNA Polymerase (New England Biolabs) was used for PCR reactions, following the manufacturer’s protocol.

### Microscopy

For scanning electron microscopy, the first fresh leaves from two-week old plants were fixed by two-side tap to observe at 5.0 kv and 3.0 pA current in Quanta 3D FEG FIB/SEM Dual Beam System.

For counting the number of nuclei per trichome initiation site, the first leaves were collected and fixed in FAA solution and stained with 4,6-diamidino-2-phenylindole (DAPI), as described previously (Walker et al., 2000). The number of nuclei per trichome initiation site (TIS) were observed with either a 10X or 20X objective under Leica DM6B fluorescent microscope. Five TIS per leaf on a total of ten leaves were examined for each line.

### Split-Luciferase Assays

Plasmids were extracted from bacterial cultures with a Midi Prep kit (Invitrogen). One ug/ul of each plasmid was introduced into Arabidopsis protoplasts derived from four-weeks old plants by polyethyene glycol-mediated transfection and incubated overnight at room temperature (Fujikura, Horiguchi, & Tsukaya, 2007; Kato & Jones, 2010). After addition of ViviREN Live Cell substrate (Promega), luminescence was measured in a Veritas microplate luminometer as described previously (Fujikura et al., 2007; Kato & Jones, 2010).

### Yeast two hybrid analysis

LiAc yeast transformation was performed as described in the GAL4 Two-Hybrid Phagemid Vector Kits manual (Agilent Technologies). The constructions including either DNA binding domain or activation domain were co-transformed into yeast strain PJ69-4a, with genotype *a trp1-901 leu2-3,112 ura3-52 his3-200 gal4Δ gal80Δ LYS2::GAL1-HIS3 GAL2-ADE2 met2::GAL7-lacZ* (James, Halladay, & Craig, 1996), and transformants were selected on SD-Leu-Trp media. For scoring interactions, cells were plated on SD-Ade-His-Leu-Trp media containing 4mM 3-Amino-1,2,4-triazole. Lac Z colony assays were conducted on filter lifts frozen in liquid nitrogen and thawed, as described in the GAL4 Two-Hybrid Phagemid Vector Kit manual (Agilent Technologies).

## Statistical methods

All statistical tests were conducted using the program Prism 8 (GraphPad Software).

### Accession numbers

The accession numbers of the main genes mentioned in this study are as follows: At5g04470 (*SIM*), At3g10525 (*SMR1/LGO*), At3g48750 (*CDKA;1*), AT3g54180 (*CDKB1;1*), AT2g38620 (*CDKB1;2*), AT2g22490 (*CYCD2;1*), AT4g34160 (*CYCD3;1*), AT5g65420 (*CYCD4;1*), AT4g03270 (*CYCD6;1*), At4g27230 (*H2A*), At5g22880 (*H2B)*, At1g68640 (*PAN*).

## Acknowledgements

This work was supported in part by a National Science Foundation award (MCB1615782) to J.C.L. and N. Kato. The authors wish to thank Hirofumi Harashima for the generous gift of plasmids containing the coding regions of *CYCD3;1* and *CYCD6;1*, Lieven De Veylder for the generous gift of plasmids containing the coding regions of *CYCD2;1* and *CYCD4;1*, John Woolford for the generous gift of yeast strain PJ69-4a, and the staff of the Socolofsky Microscopy Facility, part of the Shared Instrumentation Facility at Louisiana State University, for assistance with microscopy.

---

Supplemental Figure S1. Wild-type SIM and the Motif C mutant can both activate the *lacZ* reporter in the yeast two hybrid system. LacZ colony assays on filter lifts are shown for the indicated interaction genotypes.

